# *In Vitro* and Computational Evaluation of Thrombolytic Activity of Kinemakinase of Kinema, an Indigenous Fermented Food of Eastern Nepal

**DOI:** 10.64898/2026.08.11.744146

**Authors:** Madhu Gupta, Sumit Gupta, Mamta Kumari Gupta, Fuleshwor Mandal, Sanjay Raj Baral, Pradeep Kumar Shah, Shiv Nandan Sah

**Author notes:** **Corresponding author:** Shiv Nandan Sah.

## Abstract

Kinema is a traditional fermented soybean food indigenous to the eastern Himalayan regions of Nepal and India. The fermentation process is primarily mediated by the bacterium *Bacillus subtilis*, which produces several bioactive compounds and enzymes with potential therapeutic applications. Considering the growing burden of cardiovascular diseases and the need for effective fibrinolytic agents for thrombolytic therapy, this study aimed to extract, partially purify, and evaluate the thrombolytic potential of kinemakinase derived from kinema prepared from white soybeans. Partial purification of the enzyme was achieved using ammonium sulfate precipitation. Thrombolytic activity was assessed in vitro using human blood clots, where three enzyme dilutions demonstrated clot lysis ranging from 66% to 68%, indicating considerable fibrinolytic potential. In silico analyses were also performed to investigate the structural and functional characteristics of the enzyme. The tertiary structure obtained from UniProt was modeled using the Robetta server and refined with GalaxyRefine. Docking with fibrin using ClusPro 2.0 and molecular dynamics simulations using iMODS confirmed favorable interaction and structural stability, while disulfide engineering enhanced protein stability. The findings suggest that kinema-derived kinemakinase may serve as a promising alternative thrombolytic agent, warranting further biochemical characterization and dosage optimization.

## Introduction

Soybean-based fermented products are extensively consumed in several parts of Asia, primarily due to their unique aroma, flavor, and numerous health benefits [1]. Kinema is a conventional, non-saline fermented soybean product that holds significant cultural importance in the eastern hills of Nepal and North India [2–4]. Soybeans constitute a substantial source of phenolic compounds, including flavonol, hydroxycinnamic acid, isoflavone, rutun, daidzin, genistein, and daidzein [5, 6].These compounds transform during fermentation, resulting in a more bioactive and bioavailable form [6]. Additionally, soybean products(fermented) contain peptides that possess various properties such as antioxidant, antidiabetic, antimicrobial, antitumor, and angiotensin I converting enzyme inhibitory [7]. Traditionally, the Preparation of Kinema, a non-salted and alkaline fermented food, is done by using cooked soybeans that have undergone natural fermentation. Many bioactive components are present in fermented kinema, although research findings regarding the impact of fermentation duration on kinema’s bioactivity are limited [8].

During the fermentation process, soybeans are colonized by a diverse array of microorganisms, including bacteria, yeasts, and molds. Among these, *Bacillus* spp. is the most dominant species [9]. The predominant species identified in kinema is *Bacillus subtilis*, followed by *B. glycinifermentans*, *B. cereus*, *B. licheniformis*, *B. thermoamylovorans*, *B. coagulans*, *B. circulans*, *B. paralicheniformis*, and *Brevibacillus borstelensis* [3, 4]. Kinema is traditionally consumed by Indigenous communities in the eastern hills of Nepal [10]. It possesses a short shelf life, a slimy texture, and a strong ammonia odor. It is comparable to other fermented soybean products, including the Japanese natto with *Bacillus*, the Korean chungkukjang, the northern Thai thuanao, the northern Burmese pepock, and the Cambodian seeing [11]. The traditional preparation and consumption of kinema are prevalent among the indigenous communities of the eastern hilly regions of Nepal, Sikkim, and Darjeeling in India, as well as some regions of Bhutan. *Bacillus subtilis* is identified as the primary microorganism in the kinema through the solid-state fermentation process [2].

The use of microorganisms to synthesize bioactive compounds from fermented soybean products is effective in producing enzymes [12–14]. Studies have suggested that phenolic compounds and other bioactive components may be present in kinema, as indicated by research [15, 16]. Consuming diets high in plant phenolics over the long term may offer some protection against the development of various diseases, such as cardiovascular diseases, osteoporosis, cancers, diabetes, and neurodegenerative diseases, according to epidemiological studies and meta-analyses [8, 17]. Furthermore, these diets may improve gut microbiota and aid in body-weight management.

Nattokinase is an enzyme produced by *B. subtilis* during the fermentation of soybeans for the preparation of natto and is recognized as a potent blood–clot–dissolving protein used in the treatment of cardiovascular diseases. The enzyme demonstrates direct fibrinolytic activity by hydrolyzing fibrin and plasmin substrates, promoting the conversion of endogenous pro-urokinase into its active form, urokinase (uPA), degrading plasminogen activator inhibitor-1 (PAI-1), and increasing tissue plasminogen activator (t-PA) levels, thereby enhancing overall fibrinolysis [18].

A similar enzyme found in kinema, showing thrombolytic activity similar to Nattokinase, can be called Kinema kinase, produced by *B. subtilis*. The partially purified enzyme from freshly prepared Kinema was used in this study. A bioinformatic analysis study showed compact molecular docking between fibrin and kinemakinase (protease) enzymes. It means this enzyme can lyse blood clots, showing thrombolytic activity.

## Materials and methods

### In Vitro Study

#### Soybean Collection and Preparation of Kinema

Two kilograms of White soybeans were purchased from the Grocery of Bharatpur municipality, Chitwan, Nepal. The soybean grains were cleaned, and the remaining mud, dust, and grime were washed away with water. This study was carried out from February to March 2024 in the Biochemistry laboratory of Chitwan Medical College, Bharatpur, Chitwan, Nepal. Ethical approval was taken from the CMC-IRC (Institutional Review Committee) and lab work was performed according to Good Laboratory Practice (GLP) guidelines.

The soybean grains were soaked overnight and subsequently boiled in water for 2 h in a tightly covered container until adequate softening of the beans was achieved. To expose the cotyledons, soybean seeds were macerated by hand at around 40°C, removing the seed coat. The split seeds were maintained in a bamboo container covered with banana leaves after being carefully mixed with 1% firewood ash, and any surplus water was drained.

After being wrapped in a muslin cloth, the sample was placed in an environmental/stability cabinet room for 72 hours to undergo spontaneous fermentation at 30 ± 2 °C [10]. According to Khadka et al. (2024), the microorganisms responsible for the fermentation of kinema originate from sources such as soybeans, machinery, firewood ash, and packaging materials [19].

#### Extraction and partial purification of Kinemakinase

The enzyme was extracted from kinema that had been fermented for three days by using sodium phosphate buffer (pH 7, 50 mM) as the extraction solvent, following a kinema-to-buffer ratio of 2:3 (w/v). Specifically, 80 grams of kinema were mixed with 120 milliliters of the buffer solution. This mixture was stirred for 25 minutes with a magnetic stirrer, then filtered through clean muslin cloth to obtain a total volume of 120 milliliters. The filtrate was subsequently centrifuged at approximately 7000 rpm for 10 minutes using a centrifuge tube [19]. To ensure clarity, the centrifuged mixture was again filtered through muslin cloth using the same buffer to yield a final 120 ml enzyme extract [20].

#### Partial purification by Ammonium Sulphate

The crude extract (120 ml) of the protease was precipitated by mixing with gradually increasing concentrations of 30%, 40%, 50%, and 60%, 70%, 80% saturated ammonium sulfate at refrigerated conditions, followed by centrifugation at about 7000 rpm for 10 minutes to obtain a partially purified protease enzyme. The precipitate was centrifuged and resuspended in 50 mM sodium phosphate buffer (pH 7.0) to a final volume of 12 mL, following a previously described method with minor modifications. Total protein concentration was subsequently quantified using an ultraviolet spectrophotometer [21].

#### Confirmation of Protein in the Purified Sample

The presence of protein in the purified sample was confirmed by the development of a violet coloration in the reaction mixture by the biuret test, which is indicative of a positive protein reaction [22].

#### Determination of protein content

The protein concentration was determined by measuring the absorbance of the sample at 540 nm and comparing it with a bovine serum albumin (BSA) standard curve ranging from 0.1 to 1.2 mg/ml. The results were expressed as milligrams per milliliter [19].

#### Thrombolytic activity of protease (Blood Clot Lysis Assay)

As human blood was needed in this work, ethical approval was taken from Ethical committee of Chitwan Medical College. Venous blood samples were collected from two healthy volunteers and dispensed into sterile, pre-weighed microcentrifuge tubes (500 µL per tube). The tubes were incubated at 37 °C for 45 min to allow clot formation. Following clotting, the serum was carefully removed without disturbing the clots, and the tubes were reweighed to determine clot mass by subtracting the weight of the empty tube from that of the clot-containing tube. Each tube was properly labeled, and 100 µL of either crude or dialyzed enzyme was added to the clots. The samples were then incubated at 37 °C for 90 min to assess clot lysis [23].

For quantitative analysis, four Spinwin conical-bottom tubes (50 mL) were used, each containing 2 mL of venous blood, which was allowed to clot under controlled conditions. Crude enzyme solutions were prepared at concentrations of 100%, 75%, and 50% and added to the appropriately coded tubes containing the blood clots. Distilled water (100 µL) served as the negative control. Following incubation, the extent of clot lysis was quantified by calculating the percentage of clot lysis using the formula given below.

#### Formula for clot lysis percentage

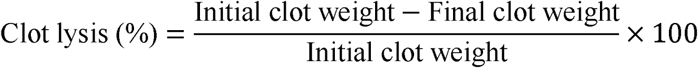

## In Silico Study

### Retrieval of the protease enzyme sequence of *Bacillus subtilis*

The amino acid sequence of the protease enzyme Subtilisin from *Bacillus subtilis* (UniProt ID: P35835) was retrieved in FASTA format from the UniProt database [24]. UniProt is a comprehensive, high-quality, and freely accessible resource that provides curated protein sequences along with detailed functional annotations, making it widely used for protein-related bioinformatics analyses [25].

### Physicochemical behavior and peptide solubility analysis

The ProtParam tool on the ExPASy server (https://web.expasy.org/protparam/) was used to evaluate the vaccine construct’s physicochemical characteristics, such as molecular weight, theoretical isoelectric point (pI), instability index, aliphatic index, amino acid composition, and grand average of hydropathicity (GRAVY) [26]. ProtParam, part of the ExPASy bioinformatics resource portal, is a widely utilized algorithm that provides a comprehensive analysis of protein sequences to determine attributes critical for protein stability, expression, and interaction [27].

### Prediction of Secondary and Tertiary Structure

The secondary structure of the protease was predicted using SOPMA [28] and PSIPRED (Buchan & Jones, 2019). The protein’s tertiary structure was initially estimated using the 3Dpro tool from SCRATCH (http://scratch.proteomics.ics.uci.edu/) [29]. Visualization of the three-dimensional structure was performed using Discovery Studio Visualizer.

To enhance the accuracy and quality of the initial model and ensure it closely resembled experimentally determined structures, the model was refined using GalaxyRefine (http://galaxy.seoklab.org/cgibin/submit.cgi?type=REFINE) [30]. This refinement process employs molecular dynamics simulations to relax the overall structure and optimize side-chain conformations (Heo et al., 2013). The refined 3D structures were further visualized using SPdbViewer and Discovery Studio [31].

The structural integrity of the tertiary models was validated by generating Ramachandran plots using PROCHECK in PDBsum (https://www.ebi.ac.uk/thorntonsrv/databases/pdbsum/Generate.html) [32], allowing assessment of residue conformations and overall stereochemical quality.

### Molecular docking analysis

Molecular docking analysis between the protease enzyme and fibrin (Fibrin 1fzc) was carried out using the ClusPro 2.0 server (https://cluspro.bu.edu/home.php) [33], a well-established and freely available web-based platform for protein–protein docking [33, 34]. The tertiary structure of fibrin (PDB ID: 1fzc) was retrieved from the Protein Data Bank (PDB) and visualized using Discovery Studio. To obtain a refined structure, non-essential components such as water molecules and ligands were removed, resulting in a cleaned PDB file [31]. The structure was subsequently processed in DeepView/Swiss-PdbViewer, where all atoms were selected and subjected to energy minimization. The energy-minimized structure was saved in PDB format and reopened in Discovery Studio, where ligands were deleted while hydrogen atoms and water molecules were retained, generating a prepared receptor file suitable for docking analysis [35]

ClusPro facilitates the docking workflow through a user-friendly interface that accepts two PDB-formatted input files. It utilizes Fast Fourier Transform (FFT)-based algorithms to sample billions of possible conformations, thereby significantly advancing protein–protein docking methodologies. Nevertheless, the accuracy of the docking results is influenced by the rigid-body assumption inherent to the approach, despite the implementation of “soft” docking strategies that permit limited structural overlap. Furthermore, the docking calculations rely on energy functions represented as correlation sums [33]. Swiss-PdbViewer v4.1 provides functionalities such as structural alignment, homology modeling, mutation analysis, and energy minimization, underscoring its utility in protein structure preparation and refinement [35].

### Molecular dynamics simulation analysis

Molecular dynamics simulations of the docked protease–fibrin complex were performed using the iMODS server (https://imods.iqf.csic.es/) [36]. Fibrin, derived from fibrinogen, a key blood plasma protein, serves as the receptor for the protease enzyme during the blood clotting process. iMODS is an advanced platform for normal-mode analysis (NMA) that facilitates exploration of protein flexibility and large-scale conformational changes. It enables the generation of transition pathways between homologous structures, even for large macromolecular complexes. The server offers features such as vibrational mode analysis, animation of molecular motions, morphing trajectories, and enhanced visualization tools, including a refined affine-model-based arrow representation to depict motion directions [35].

### Disulfide engineering

Using the Design v2.0 web server, additional disulfide bonds were introduced into the newly built construct to increase its structural stability [37]. The three-dimensional structure of Subtilisin was initially assessed using the ProtParam tool [26] to evaluate its physicochemical properties. The disulfide engineering algorithm employed by Design v2.0 integrates structural geometry analysis, energy minimization, and molecular modeling to identify and validate potential disulfide bond sites. This strategy aims to reinforce protein stability by promoting correct folding, enhancing thermostability, and increasing resistance to denaturation through the strategic incorporation of disulfide bridges [37].

### Codon optimization in silico cloning (Expression analysis)

The coding DNA sequence (cDNA) of kinemakinase was reverse-translated from its corresponding amino acid sequence using the EMBOSS Backtranseq tool (https://www.ebi.ac.uk/jdispatcher/st/emboss_backtranseq). During this process, each amino acid was converted into its appropriate nucleotide triplets based on a standard codon usage table, in which 64 codons encode 20 amino acids and translational stop signals [38]. To enhance heterologous expression efficiency, the translated gene sequence was subsequently optimized using the Java Codon Adaptation Tool (JCAT) for *Escherichia coli* (strain K-12) as the expression host [39]. Codon optimization was performed by adjusting codon usage to match host preferences while evaluating key parameters such as the Codon Adaptation Index (CAI) and GC content. In addition, sequence elements detrimental to efficient expression, including prokaryotic ribosome-binding sites, Rho-independent transcription terminators, and restriction enzyme recognition sites, were systematically excluded [40]. The codon-optimized gene was then in silico cloned into the *E. coli* pET-28a(+) expression vector using the SnapGene software to verify correct insertion and to simulate successful expression of the optimized construct [41]

## Results

### In vitro work

In this study, Kinema was prepared in the laboratory following the traditional methods. The fermentation process was over in 3 days, producing a soft, sticky (Slimy appearance), and ammonical, pungent kinema with a distinct flavor and aroma. It was slightly alkaline in taste (Figure 1C).

**Figure 1:**
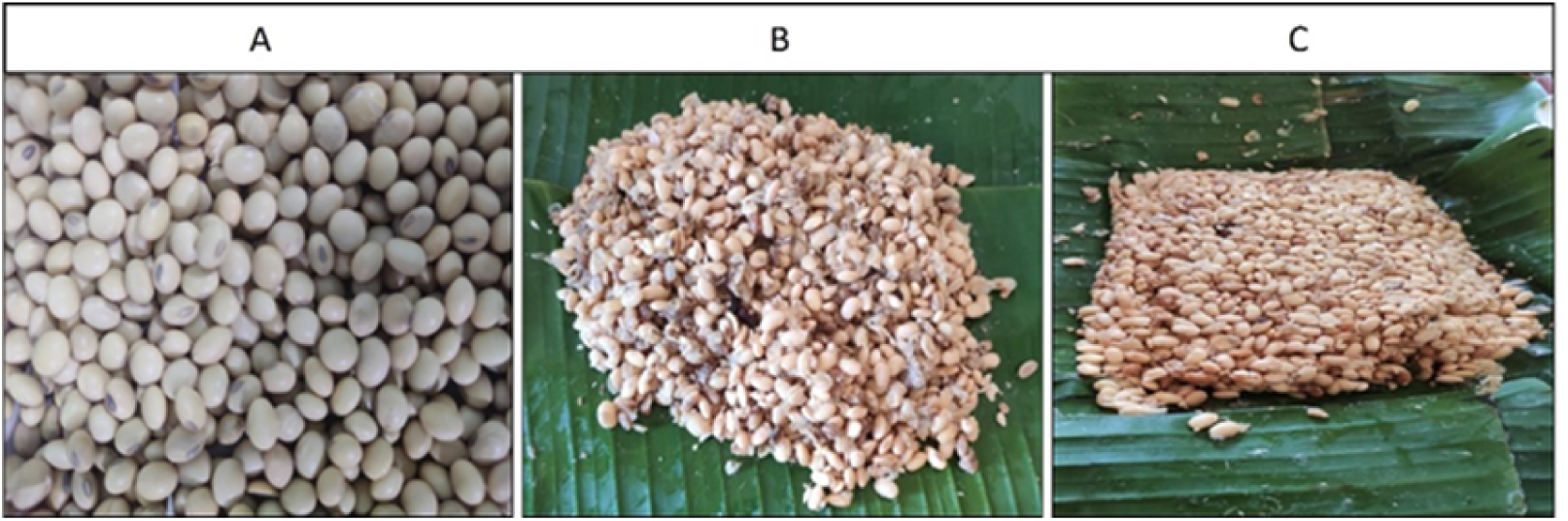
Process of Kinema preparation. A. Soyabean B. Cooked soybeans on a banana leaf (ready to ferment) C. Prepared Kinema

### Extraction and partial purification of the enzyme

By mixing kinema into phosphate buffer (2:3), the crude protease enzyme from kinema was successfully extracted. Partial purification was achieved by using ammonium sulfate by increasing its concentration up to 80%. Precipitated protein was dissolved in phosphate buffer.

### Evaluation of partially purified protease enzyme

A positive biuret test showed the protein content of a partially purified enzyme. An enough amount of Protease content was found in the partially purified enzyme content. At this step, total enzyme content was measured in an Ultraviolet (UV) spectrophotometer (Figure 2).

**Figure 2:**
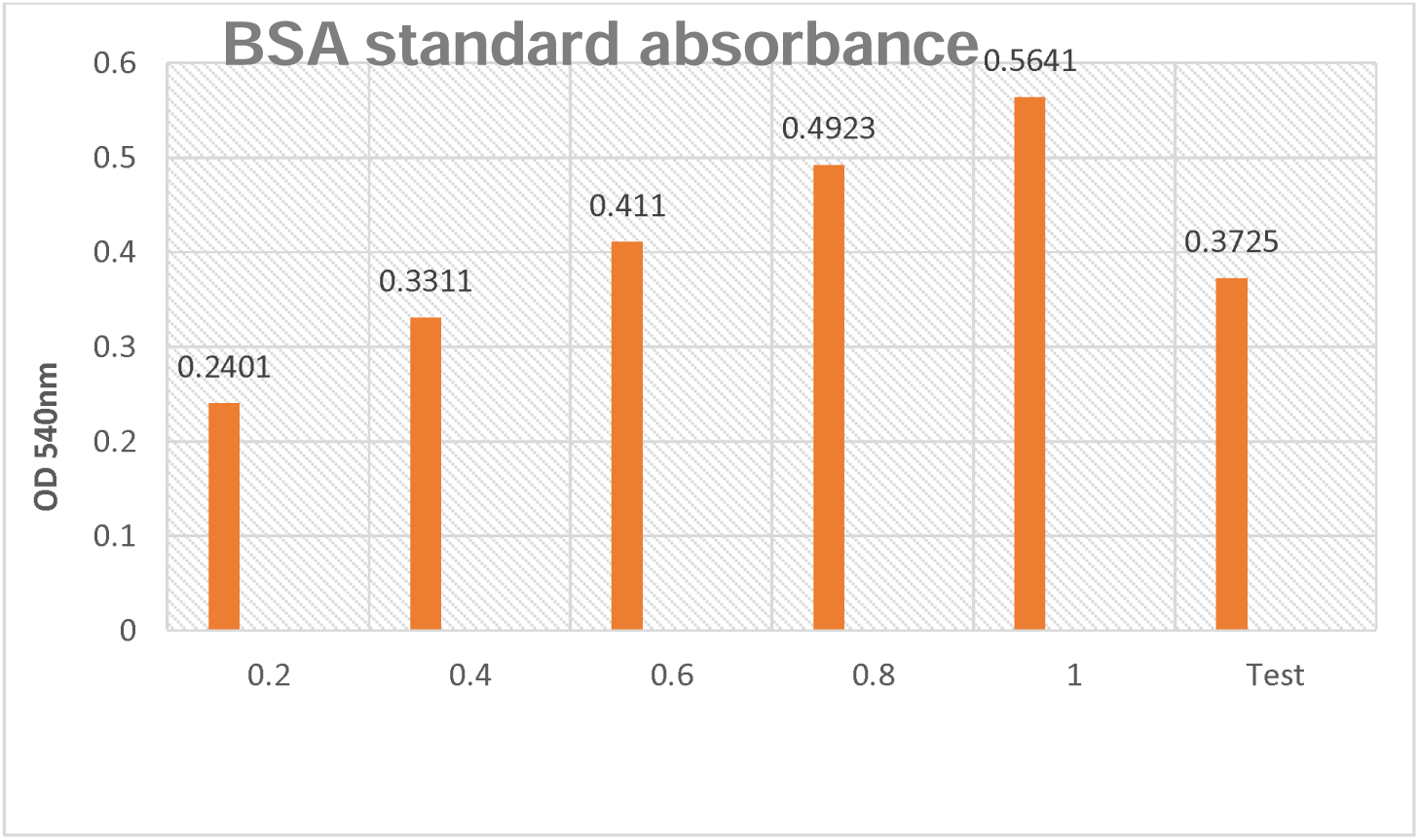
BSA standard absorbance. Partially purified kinemakinase OD was 0.3725; the calculated protease content was found to be 5 mg/ml.

**Figure 3:**
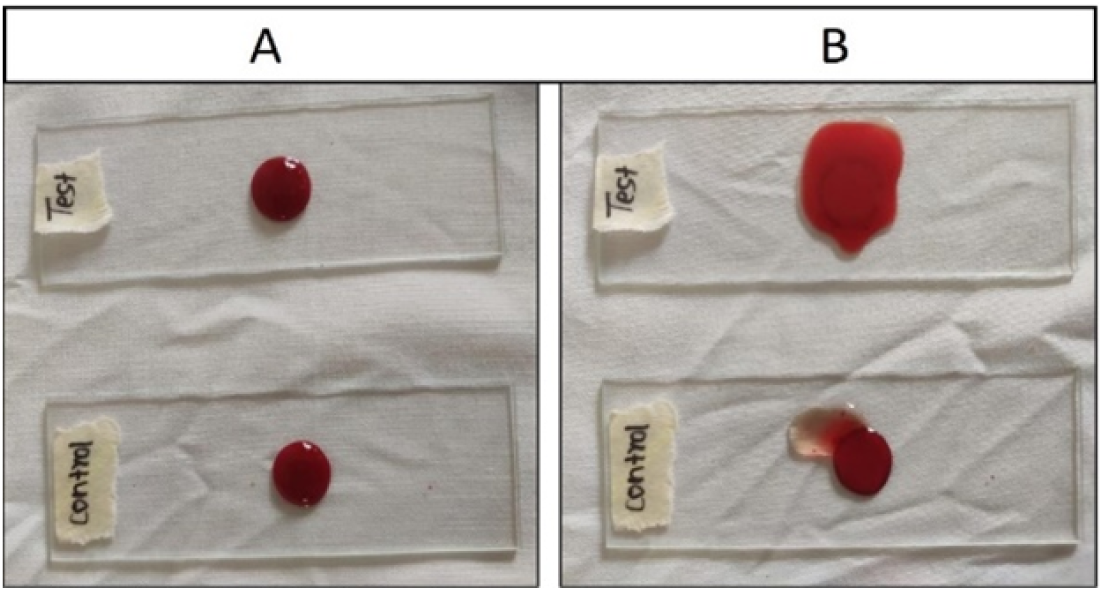
Blood clot lysis (qualitative) A. Test and control blood clots (before adding kinemakinase and sterile distilled water, respectively) B. Positive lysis in the Test and negative lysis in the control blood clot (after adding kinema kinase and sterile distilled water, respectively)

### Evaluation of blood clot lysis (thrombolytic activity)

Qualitative evaluation of blood clot lysis with kinema kinase extracted from kinema on the first glass slide (test), compared with sterile distilled water on the second glass slide (control), was done. The results demonstrated effective clot lysis upon treatment with kinema extract on blood clots derived from healthy individuals (Figure 4). Three different dilutions of the extract, 100%, 75%, and 50%, were applied, resulting in clot lysis percentages of 68.001%, 67.011%, and 66.101%, respectively (Figure 5). Each of these values showed a statistically significant difference compared to the control group treated with water. Overall, the clot lysis percentage induced by various concentrations of kinema extract ranged from 66.101% to 68.001%.

**Figure 4:**
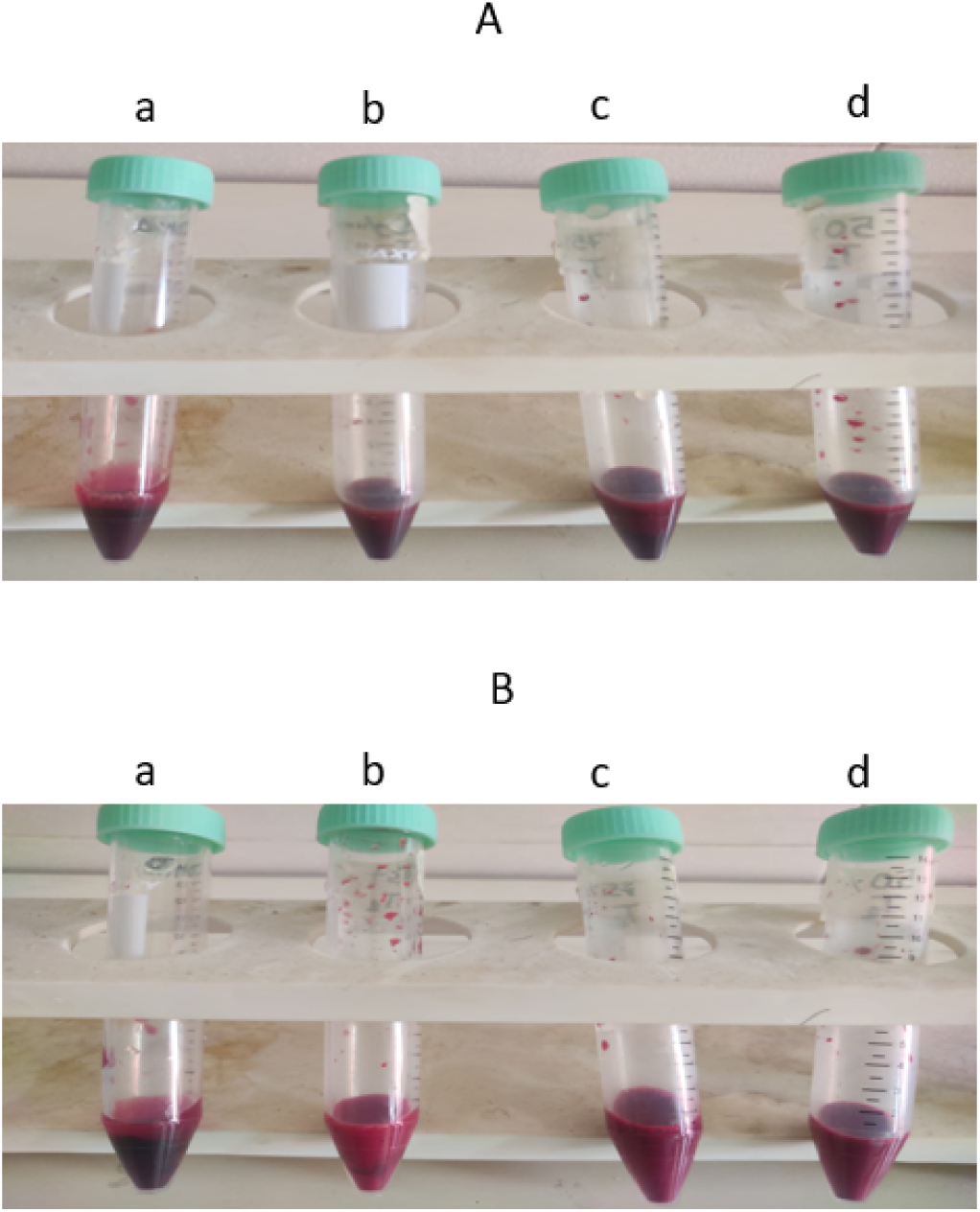
Blood Clot Lysis (quantitative) A. Before clot lysis (a, b, c, and d) B. After Clot lysis a. Blood clot + water b. Blood clot +100% kinemakinase (partially purified) c. Blood clot + 75% kinemakinase (partially purified) d. Blood clot +50% kinemakinase (partially purified)

**Figure 5:**
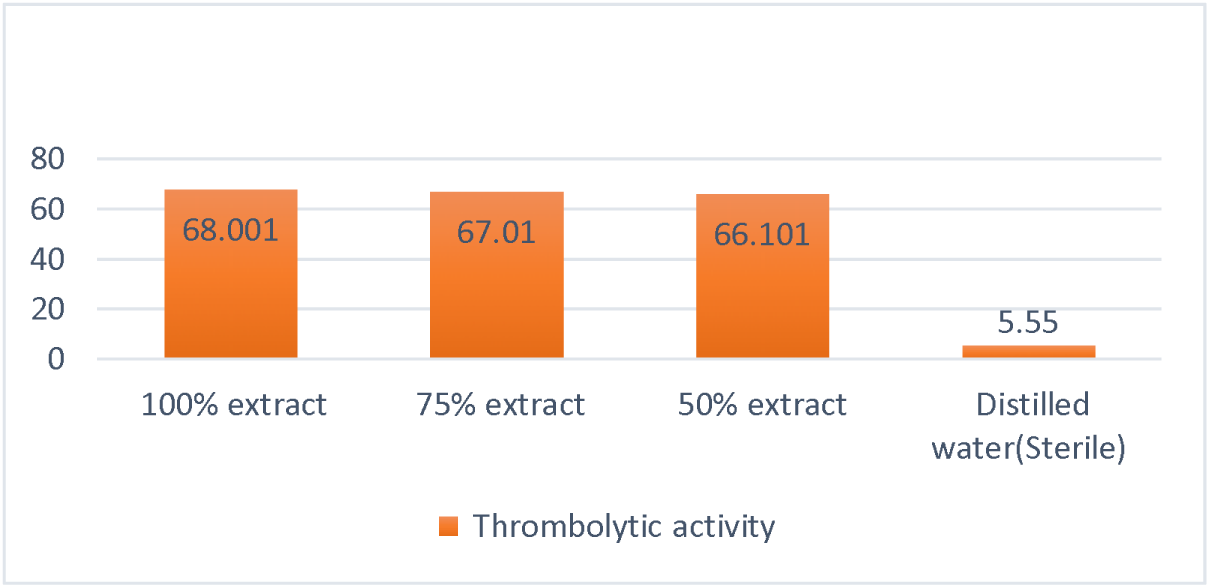
Thrombolytic activity of kinema kinase (Clot lysis of blood sample of a normal subject by different concentrations of kinema extract)

## Bioinformatics analysis

### Retrieval of the protease enzyme sequence of *Bacillus subtilis*

The protease enzyme sequence of kinema kinase, comprising 381 amino acids with an estimated molecular weight of approximately 39.5 kDa, was retrieved from the UniProt database. The theoretical isoelectric point (pI) of the protein was calculated to be 9.04. Physicochemical analysis revealed an instability index of 28.84, classifying the protein as stable. The aliphatic index was determined to be 81.73, indicating high thermostability, while the grand average of hydropathicity (GRAVY) value of −0.055 suggested a slightly hydrophilic nature. A detailed summary of the physicochemical properties of kinemakinase is provided in Supplementary Table S1.

Secondary structure prediction of the kinema kinase protein was performed using the SOPMA server, which indicated that the final construct consisted of 27.30% α-helices, 19.95% extended strands, and 57.76% random coils (Figure 6B). For tertiary structure prediction, the 3Dpro tool from the SCRATCH suite was employed, and the predicted model was subsequently refined using the GalaxyRefine server to improve structural accuracy and stability. GalaxyRefine generated five refined models, among which Model 1 was selected for further analyses based on overall quality assessment (Supplementary Table S2).

**Figure 6:**
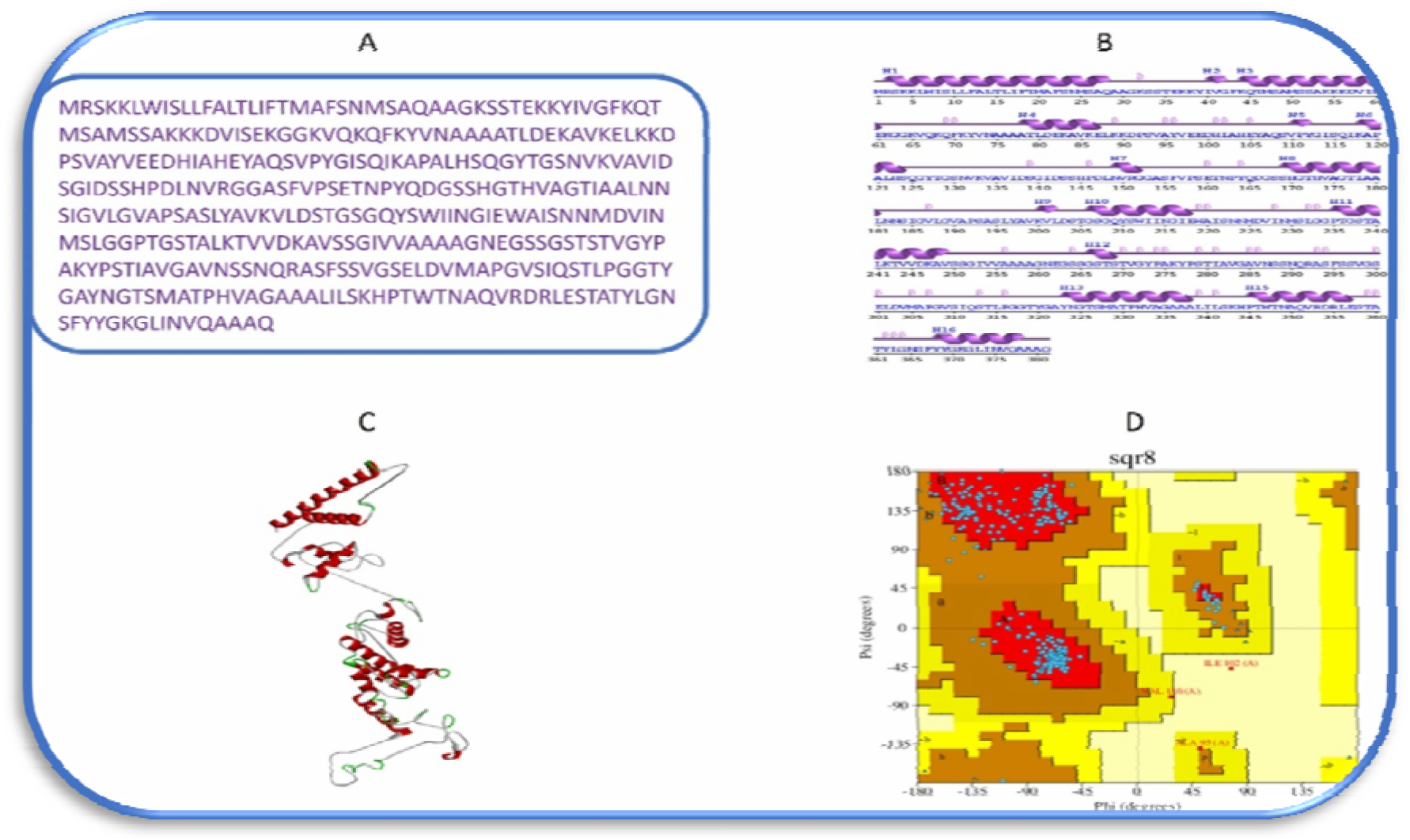
Schematic presentation of finalized Kinema Kinase construct: A) Kinema-Kinase sequence B) Secondary structure of Kinema-kinase C) Tertiary structure of Kinema-Kinase D) Validation of tertiary structure of Kinema-kinase by Ramachandran Plot using PROCHECK

**Figure 7:**
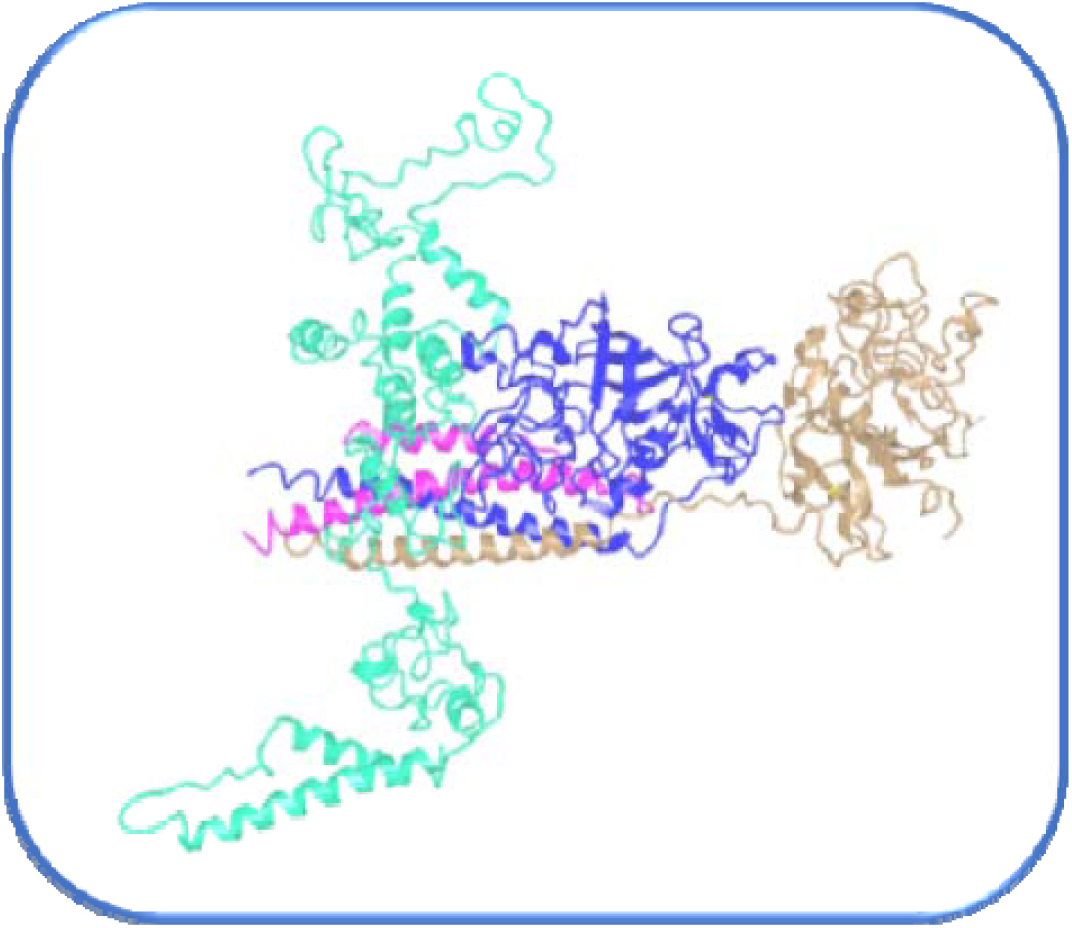
Schematic representation of the Kinemakinase-Fibrin docked complex (the construct is depicted in cyan, the receptor in green, and the complex itself in red-purple)

Structural validation of Model 1 demonstrated high reliability, with a root mean square deviation (RMSD) of 0.381 Å, a MolProbity score of 1.8, a clash score of 12.4, and no detected bad rotamers. Ramachandran plot analysis revealed that 96.8% of residues were located within favored regions. Further validation using PROCHECK showed that 302 residues (92.1%) were present in the most favored regions, 23 residues (7.0%) in additionally allowed regions, 2 residues (0.6%) in generously allowed regions, and only 1 residue (0.3%) in the disallowed region. Overall, 99.7% of residues were located within allowed regions, confirming the structural stability and integrity of the refined three-dimensional model (**Figure 6D**).

### Molecular docking analysis

The ClusPro 2.0 server was utilized for molecular docking of the designed kinemakinase with host blood cell receptor fibrin (PDB ID: 1FZC) [33]. ClusPro generated 30 docked complex models, among which the top five, based on energy scores and cluster sizes, were shortlisted and presented in Supplementary Table S3. Model 1 was selected for further analysis, as it demonstrated the most favorable interaction profile, characterized by the lowest binding energy and the largest cluster size, specifically, a binding energy of -1150 kcal/mol and a cluster size of 89. These results indicate a strong and stable interaction between kinema kinase and fibrin.

To further investigate the molecular interface of the complex, the PDBsum server was employed. This tool provided a comprehensive overview of the interacting residues between kinemakinase and fibrin, along with a detailed schematic illustration of the non-covalent interactions stabilizing the docked complex [42].

### Molecular dynamics simulation analysis

The docked complex between kinemakinase and fibrin was analyzed using molecular dynamics simulation on the iMODS server [36]. This simulation assessed the structural stability of kinema kinase within the docking complex, demonstrating improved molecular flexibility and stability. The kinemakinase-fibrin complex was evaluated using multiple analytical approaches, including three-dimensional structural modeling, B-factor (atomic mobility) analysis, eigenvalue assessment, variance analysis, and covariance mapping to elucidate the dynamic behavior and stability of the complex (Figure 8).

**Figure 8:**
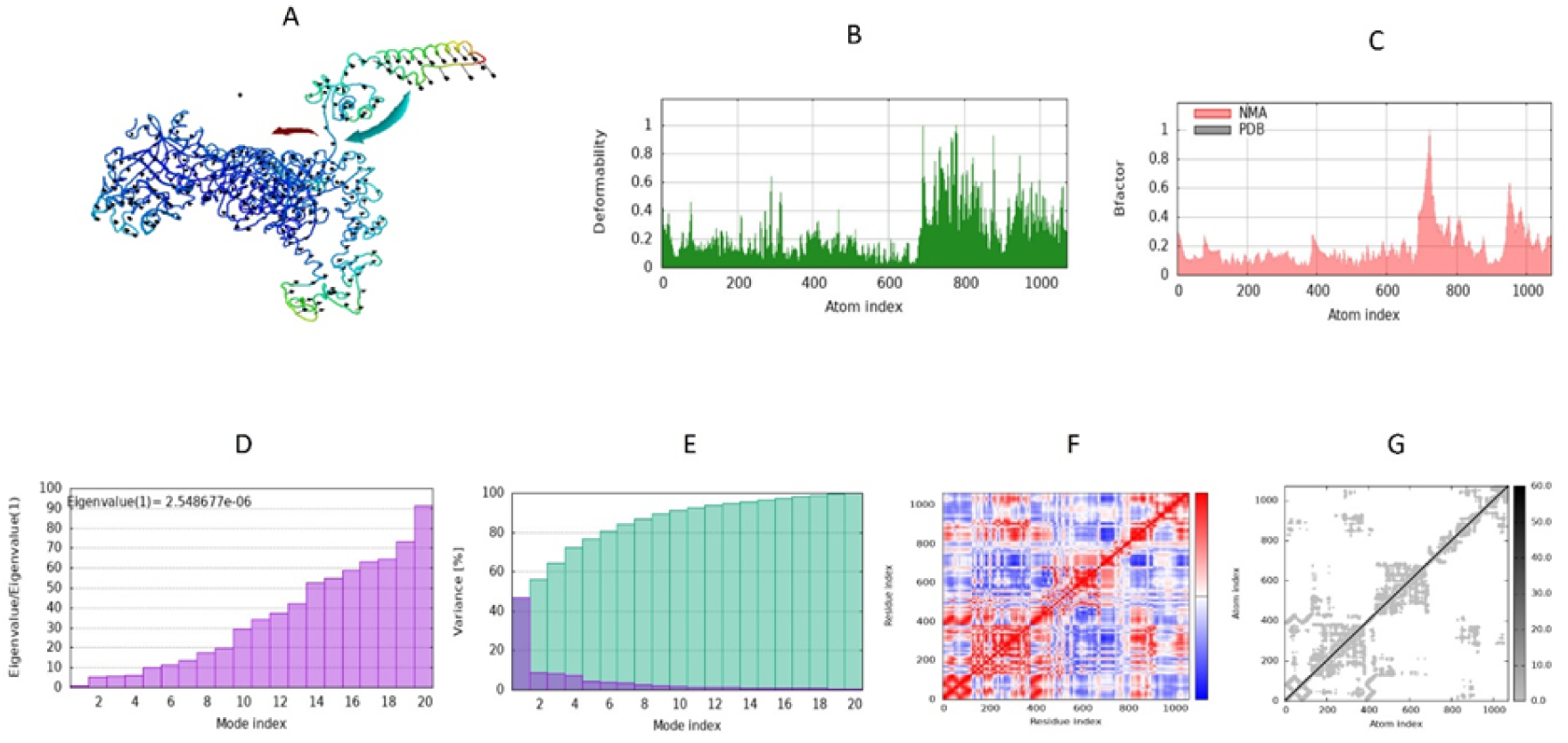
Molecular dynamics simulations of the Fibrin-Kinemakinase complex. (A) Fibrin-Kinema kinase docking complex (3D structure) with NMA mobility (B) Deformability (C) B-factor (D) Eigenvalue (E) Variance (F) Covariance map (G) Elastic Network

Protein motion was further analyzed using main-chain deformability graphs (Figure 8B), which highlighted peaks indicating areas of high deformability, particularly in the hinge regions. The B-factor values provided insight into the flexibility of the kinemakinase candidate by measuring the variability of each atom.

The association between the PDB structure for the docked complex and normal mode analysis (NMA) was demonstrated by the B-factor graph.

Elastic network models (ENMs) were constructed for the kinema kinase–fibrin complex to visualize atomic interactions represented as pairs of atoms interconnected by virtual “springs” (Figure 8A). In this model, each dot corresponds to a spring between an atomic pair, with the color intensity reflecting the rigidity of the interaction; darker gray dots indicate stiffer springs and reduced flexibility. The covariance matrix analysis (Figure 8F) further illustrated residue–residue motion correlations within the complex, where red denotes positively correlated movements, white indicates the absence of correlation, and blue represents anti-correlated motions. Additionally, the B-factor profile closely mirrored the root-mean-square fluctuation (RMSF) values (Figure 8C), confirming consistency between atomic mobility and structural flexibility assessments. Overall, the ENM approach provided a simplified yet effective representation of the intrinsic dynamics of the docked complex, facilitating deeper insight into large-scale macromolecular motions and stability.

### Disulfide engineering

The Disulfide by Design v2.0 web server was employed to introduce multiple disulfide bonds into the three-dimensional structure of kinema kinase to enhance its structural stability [37]. This method is perfect for protein engineering because it enhances protein stability by promoting covalent connections between disulfide bonds that adhere to particular geometric conformations. A new disulfide engineering method can create these disulfide links in target proteins [43, 44].

The Disulfide by Design (DbD) 2.12 server was used to introduce stabilizing disulfide bonds into the kinemakinase structure [37]. The GalaxyRefine-optimized PDB file of the kinema kinase three-dimensional structure was uploaded to the server, where residue pairs were systematically evaluated for potential cysteine substitutions capable of forming energetically favorable disulfide linkages based on geometric and energetic criteria [45]. Nineteen of the 36 latent amino acid pairs that were produced were chosen for disulfide engineering to modify cysteine residues. These particular amino acid pairs were given new disulfide links, which improved kinemakinase stability without changing the PDB structure as a whole (Figure 9). Supplementary Table S4 contains a list of the amino acid pairings that were chosen for mutation.

**Figure 9:**
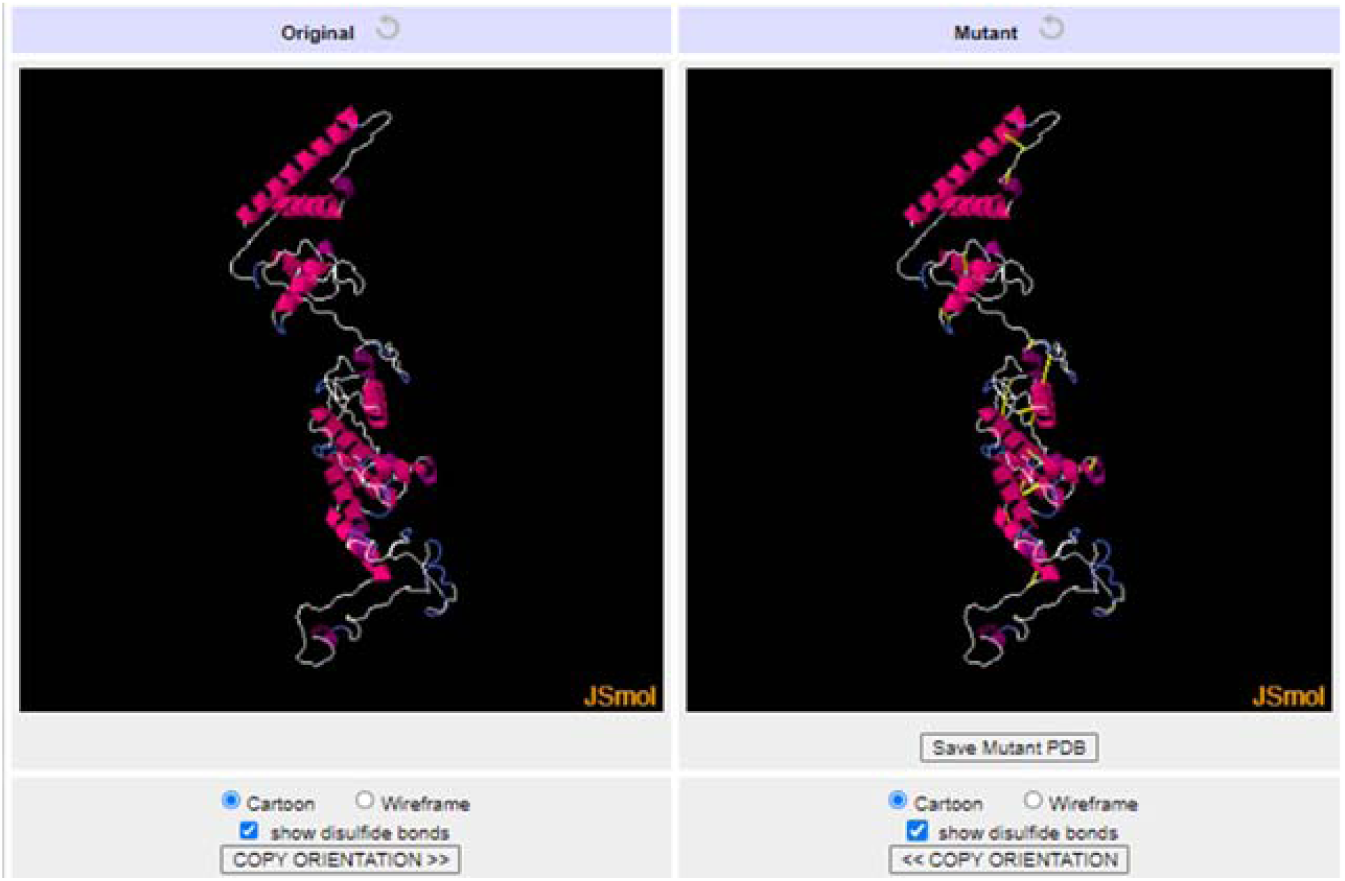
Disulfide engineering.

## Discussion

Kinema is a product that can be prepared through the process of fermenting partially cooked soybeans on banana leaves. This procedure involves the use of *B. subtilis*, a critical fermenting bacterium that produces various enzymes and beneficial bioactive substances that contribute positively to human health.

*Bacillus subtilis* is a bacterium that produces a unique protease enzyme during the fermentation process, which breaks down the soy protein on the surface of white soybeans [3]. The slimy and sticky texture of Kinema is attributed to the mucilage produced by soybean extracellular enzymes, which contain exopolypeptides of D-isomeric glutamic acid. The proportion of nitrogen soluble in water and trichloroacetic acid increases while the protein nitrogen content decreases significantly [46]. Proteases produced by *Bacillus subtilis* hydrolyze soybean proteins into polypeptides during fermentation. It has been reported that kinema fermentation leads to a substantial increase in free amino acids and ammonia levels, by approximately 40-fold and 60-fold, respectively [47]. Furthermore, the fermentation process imparts a characteristic slimy texture and a strong ammoniacal odor, both of which are considered key indicators of good-quality kinema [2].

The appeal of fermented soybean products is largely attributed to their rich content of bioactive constituents, including polyphenols such as isoflavones, phenolic acids, and flavonols, along with bioactive peptides, γ-aminobutyric acid, antimicrobial compounds, fibrinolytic enzymes, vitamins, and exopolysaccharides, as reported by Sanjukta and Rai (2016), Sharma et al., 2021 and Peiretti et al. (2019). These components collectively contribute to the nutritional, functional, and therapeutic properties of fermented soy{Citation}bean foods [5, 6, 48, 49].

*Bacillus* species possess considerable industrial significance, as they underpin numerous bioprocesses and produce a wide range of metabolites with strong commercialization potential. Their widespread application in solid-state fermentation is primarily due to their capacity to synthesize diverse hydrolytic enzymes, including cellulase, α-amylase, proteases, pectinase, and β-glucosidase, which play essential roles in substrate degradation and value-added product formation [12, 14].

Additionally, fermented soybean products have been reported to contain peptides exhibiting antioxidant, anticancer, antidiabetic, antibacterial, and angiotensin I–converting enzyme (ACE) inhibitory activities [7]. In the present study, sodium phosphate buffer (50 mM, pH 7) was employed as a solvent for the extraction of protease enzymes from freshly prepared kinema, maintaining a kinema-to-solvent ratio of 1:1.5 (w/v). Protease activity during soybean fermentation was observed to peak on the third day, followed by a gradual decline thereafter [19].

Following the partial purification of kinemakinase (Kinema protease) through ammonium sulfate precipitation, the enzyme was utilized to assess its thrombolytic activity. The enzyme has garnered significant interest due to its potential health benefits, which include promoting healthy blood flow and potentially preventing cardiovascular conditions such as heart attacks and strokes.

Researchers from Nepal, including Khadka et al. (2024), indicated that ammonium sulfate precipitates obtained from crude extracts were subjected to dialysis for partial purification [19].

Soybean, a yellow legume native to China, serves as a dietary staple for both humans and animals across many East Asian countries and is considered a cost-effective source of nutrition due to its high protein and bioactive compound content [50]. Soybeans contain approximately 37.69% protein, 28.2% crude fat, 4.29% ash, 8.07% moisture, 5.44% crude fiber, and 16.31% carbohydrates. They are particularly valued for their richness in essential amino acids, which the human body cannot synthesize and must therefore obtain through dietary intake.

In addition to their nutritional profile, soybeans offer multiple health benefits, as demonstrated by numerous clinical studies. Consumption of soy and soy-derived products has been associated with reductions in blood cholesterol levels, cardiovascular disease risk, obesity, cancer incidence, diabetes, kidney disease, and osteoporosis [51].

A study conducted by Pahadi (2022) reported an absorbance of 0.377 at 540 nm for a partially purified enzyme, which is comparable to the results of our study, which recorded an absorbance of 0.3275 at the same wavelength. This similarity in findings supports the reliability of our results [52]. Similarly, Enechi and Emilia (2013) demonstrated that the optical density measured using the biuret reagent decreased from 1.269 to 0.144 as the concentration of protein in the solution declined. Their study utilized bovine serum albumin (BSA) prepared at concentrations ranging from approximately 0.1 to 8 mg/mL, illustrating the direct relationship between protein concentration and absorbance in biuret-based protein quantification. Protein concentrations were determined using the proposed biuret method. The concentration of each unknown protein sample was then calculated based on the standard curve, taking into account the appropriate dilution factor [53], which is comparable to our findings [53, 54].

In our study, we evaluated the clot lysis (Thrombolysis) percentage using different concentrations of kinema extracts. The results showed a range of clot lysis from 66.1011% to 68.0010%. These findings are comparable to those reported by Prasad et al. (2006) and Chandrasekaran et al. (2015), who also observed significant clot-dissolving activities in their respective studies on similar subjects. The thrombolytic effect is due to the partially purified kinema protease produced by *B. subtilis*, which effectively breaks down fibrin, the main protein in clot formation [23, 55].

Selecting a suitable substrate is a key factor in enhancing enzyme production. In this study, kinema was used for protease extraction. Kinema kinase enzyme showed fibrinolytic activity similar to the study conducted by Weng et al. (2017), and nattokinase enzyme was produced in Natto. Hence, our study strongly recommends that this enzyme be named kinemakinase. Hence, the Kinemakinase enzyme in kinema is similar to the nittokinase enzyme in Natto, produced by the same bacteria, *B. subtilis.* Soybean as a substrate and *B. subtilis* are common in both cases [56].

Nattokinase is one of the most extensively studied and commercially utilized fibrinolytic proteases derived from soybean-fermented foods. It is primarily employed in therapeutic applications for the treatment of thrombotic disorders and thrombo-vascular diseases [57]. Fibrinolytic serine proteases from fermented foods are generally neutral to alkaline in nature, exhibiting optimal activity at a pH between 8 and 10 and temperatures ranging from 30 to 70 °C. In contrast, metalloproteases demonstrate peak activity within a pH range of 6 to 7 and a temperature range of 33 to 50 °C [58]. With rising incidence rates, cardiovascular diseases (CVDs) continue to be the world’s leading cause of premature mortality and disability. Common risk factors contributing to CVDs include hyperlipidemia, hypertension, diabetes, obesity, smoking, and physical inactivity [59].

All of these drugs have significant drawbacks, such as limited effectiveness, bleeding tendencies, lack of specificity, poor efficacy at low dosages, and inadequate fibrin specificity [60]. Kinema, a source of protease enzymes, has the potential to control CVDs with its antithrombotic activity and by providing essential nutrients to the body. Kinema, such as Natto, may be the ideal option for Nepalese individuals to prevent and treat CVDs through dietary means, including the consumption of kinema. As a result, there is a pressing need for safer and more cost-effective thrombolytic agents. Since Natto is an oral antithrombotic agent for the prevention of CVDs, kinema can be used as an antithrombotic agent similar to Natto. Therefore, this study is of great significance in the field of pharmacology, particularly in the context of nutraceuticals (food as medicine). By incorporating kinema into their diet, individuals can control and treat CVDs without relying on medication, saving money and reducing the prevalence of the disease. Furthermore, the increased demand for kinema in the market could create new job opportunities in the country. This study aims to evaluate and clarify the antithrombotic effects of the protease extracted from kinema.

Nattokinase enhances fibrinolytic activity through multiple mechanisms, including the direct hydrolysis of fibrin and plasmin substrates, conversion of pro-urokinase into active urokinase (uPA), degradation of plasminogen activator inhibitor-1 (PAI-1), and upregulation of tissue plasminogen activator (t-PA) levels [18]. Similarly, kinema kinase exerts its thrombolytic effect through comparable fibrinolytic pathways.

Docking studies of the engineered Kinemakinase construct with fibrin demonstrated a strong binding affinity, a key factor in facilitating fibrin hydrolysis [61]. This docking approach was instrumental in confirming the functional capability of the Kinemakinase enzyme [62]. To further substantiate these findings, molecular dynamics (MD) simulations were conducted, since docked complexes can lose stability over time, potentially impairing their fibrin-hydrolyzing function. The binding energy values showed a direct correlation with the enzyme’s functional activity and efficiency [63]. MD simulation results indicated that the complexes maintained considerable binding stability, with only minor fluctuations throughout the simulation period.

The iMODS service, which uses internal coordinate models for protein structures to perform normal mode analysis (NMA), was used to analyze the docked complex further [36, 64]. The findings revealed that the Kinemakinase-fibrin complex remained structurally stable, with any flexibility or deformability restricted to specific predicted regions, mainly at the molecular termini [31]. The deformability energy was evaluated using eigenvalues, with lower values indicating greater flexibility in the complex [65].

The interaction of proteins with other biological macromolecules depends heavily on structural flexibility [65], which was assessed through iMODS-based NMA. This technique analyzes the movement and flexibility of the docked complexes based on their atomic coordinates [36]. The presence of darker-gray dots in the analysis indicated rigid atoms with limited mobility, reinforcing the structural stability of the complex. The foundation of NMA is the idea that the most important and functionally significant molecular movements are represented by the lowest frequency normal modes [66]. The analysis identified considerable molecular mobility, highlighting the proteins’ inherent structural flexibility. This observed mobility in the NMA results further validated the dynamic adaptability of the docked complexes.

This research focused on creating an *in vitro* blood clot lysis model using Kinemakinase, a protease derived from Kinema, as a thrombolytic agent. The findings confirmed that Kinemakinase exhibits strong clot-dissolving activity, highlighting its potential for use in treating cardiovascular diseases.

## Conclusion

Kinema is one of the most nutritious and healthy fermented foods in Nepal. The major fermented bacterium is *B. subtilis*. The antithrombotic activity of kinema is highest after three days of fermentation due to bacterial protease on the soybean protein substrate. The kinema preparation process is simple and cost-effective. Protease (kinemakinase) from kinema has demonstrated potential thrombolytic activity. As a result, kinema kinase may have antithrombotic benefits and can be used as preventive medicine, but it should not be considered a substitute for standard medical treatments for conditions like cardiovascular disease or stroke.

## Acknowledgements

We would like to thank the Department of Pharmacy, Chitwan Medical College, Bharatpur, Tribhuvan University, Nepal, for providing laboratory facilities to complete this research work.

## References

1. Sourabh A, Rai AK, Chauhan A, et al (2015) Health related issues and indigenous fermented products. Indig Fermented Foods South Asia 7:309

2. Khadka DB, Lama JP (2020) Traditional fermented food of Nepal and their nutritional and nutraceutical potential. In: Nutritional and Health Aspects of Food in South Asian Countries. Elsevier, pp 165–194

3. Kharnaior P, Tamang JP (2022) Metagenomic-Metabolomic Mining of Kinema, a Naturally Fermented Soybean Food of the Eastern Himalayas. Front Microbiol 13:868383. 10.3389/fmicb.2022.868383

4. Kumar J, Sharma N, Kaushal G, et al (2019) Metagenomic Insights Into the Taxonomic and Functional Features of Kinema, a Traditional Fermented Soybean Product of Sikkim Himalaya. Front Microbiol 10:1744. 10.3389/fmicb.2019.01744

5. Peiretti PG, Karamać M, Janiak M, et al (2019) Phenolic Composition and Antioxidant Activities of Soybean (Glycine max (L.) Merr.) Plant during Growth Cycle. Agronomy 9:153. 10.3390/agronomy9030153

6. Rai AK, Sanjukta S, Chourasia R, et al (2017) Production of bioactive hydrolysate using protease, β-glucosidase and α-amylase of Bacillus spp. isolated from kinema. Bioresour Technol 235:358–365. 10.1016/j.biortech.2017.03.139

7. Sanjukta S, Rai AK (2016) Production of bioactive peptides during soybean fermentation and their potential health benefits. Trends Food Sci Technol 50:1–10. 10.1016/j.tifs.2016.01.010

8. Katuwal N, Raya B, Dangol R, et al (2023) Effects of fermentation time on the bioactive constituents of Kinema, a traditional fermented food of Nepal. Heliyon 9:e14727. 10.1016/j.heliyon.2023.e14727

9. Tamang JP, Thapa N, Bhalla TC, Savitri (2016) Ethnic Fermented Foods and Beverages of India. In: Tamang JP (ed) Ethnic Fermented Foods and Alcoholic Beverages of Asia. Springer India, New Delhi, pp 17–72

10. Tamang JP (2015) Naturally fermented ethnic soybean foods of India. J Ethn Foods 2:8–17. 10.1016/j.jef.2015.02.003

11. Tamang JP (2010) Diversity of fermented foods. Fermented Foods Beverages World 1:41–83

12. Lin H-TV, Lu W-J, Tsai G-J, et al (2016) Enhanced anti-inflammatory activity of brown seaweed Laminaria japonica by fermentation using Bacillus subtilis. Process Biochem 51:1945– 1953. 10.1016/j.procbio.2016.08.024

13. Rai AK, Kumari R, Sanjukta S, Sahoo D (2016) Production of bioactive protein hydrolysate using the yeasts isolated from soft chhurpi. Bioresour Technol 219:239–245. 10.1016/j.biortech.2016.07.129

14. Salim AA, Grbavčić S, Šekuljica N, et al (2017) Production of enzymes by a newly isolated Bacillus sp. TMF-1 in solid state fermentation on agricultural by-products: The evaluation of substrate pretreatment methods. Bioresour Technol 228:193–200. 10.1016/j.biortech.2016.12.081

15. Moktan B, Saha J, Sarkar PK (2008) Antioxidant activities of soybean as affected by Bacillus-fermentation to kinema. Food Res Int 41:586–593. 10.1016/j.foodres.2008.04.003

16. Samruan W, Gasaluck P, Oonsivilai R (2014) Total Phenolic and Flavonoid Contents of Soybean Fermentation by *Bacillus subtilis* SB-MYP-1. Adv Mater Res 931–932:1587–1591. 10.4028/www.scientific.net/AMR.931-932.1587

17. Yen G-C, Cheng H-L, Lin L-Y, et al (2020) The potential role of phenolic compounds on modulating gut microbiota in obesity. J Food Drug Anal 28:195–205. 10.38212/2224-6614.1054

18. Yatagai C, Maruyama M, Kawahara T, Sumi H (2007) Nattokinase-Promoted Tissue Plasminogen Activator Release from Human Cells. Pathophysiol Haemost Thromb 36:227–232. 10.1159/000252817

19. Khadka DB, Pahadi T, Aryal S, Karki DB (2024) Partial purification and characterization of protease extracted from kinema. Heliyon 10:e27173. 10.1016/j.heliyon.2024.e27173

20. Nafi A, Foo H, Jamilah B, Ghazali HM (2013) Properties of proteolytic enzyme from ginger (Zingiber officinale Roscoe). Int Food Res J 20:

21. Ueda M, Kubo T, Miyatake K, Nakamura T (2007) Purification and characterization of fibrinolytic alkaline protease from Fusarium sp. BLB. Appl Microbiol Biotechnol 74:331–338. 10.1007/s00253-006-0621-1

22. Buzanovskii VA (2017) Determination of proteins in blood. Part 1: Determination of total protein and albumin. Rev J Chem 7:79–124. 10.1134/S2079978017010010

23. Chandrasekaran SD, Vaithilingam M, Shanker R, et al (2015) Exploring the In Vitro Thrombolytic Activity of Nattokinase From a New Strain Pseudomonas aeruginosa CMSS. Jundishapur J Microbiol 8:. 10.5812/jjm.23567

24. Touhidinia M, Sefid F, Bidakhavidi M (2021) Design of a Multi-epitope Vaccine Against Acinetobacter baumannii Using Immunoinformatics Approach. Int J Pept Res Ther 27:2417– 2437. 10.1007/s10989-021-10262-4

25. Coudert E, Gehant S, De Castro E, et al (2023) Annotation of biologically relevant ligands in UniProtKB using ChEBI. Bioinformatics 39:btac793. 10.1093/bioinformatics/btac793

26. ProtParam E (2017) ExPASy-ProtParam tool. SIB Lausanne Switz

27. Gasteiger E, Hoogland C, Gattiker A, et al (2005) Protein Identification and Analysis Tools on the ExPASy Server. In: Walker JM (ed) The Proteomics Protocols Handbook. Humana Press, Totowa, NJ, pp 571–607

28. Deléage G (2017) ALIGNSEC: viewing protein secondary structure predictions within large multiple sequence alignments. Bioinformatics 33:3991–3992. 10.1093/bioinformatics/btx521

29. Cheng J, Randall AZ, Sweredoski MJ, Baldi P (2005) SCRATCH: a protein structure and structural feature prediction server. Nucleic Acids Res 33:W72–W76. 10.1093/nar/gki396

30. Heo L, Park H, Seok C (2013) GalaxyRefine: protein structure refinement driven by side-chain repacking. Nucleic Acids Res 41:W384–W388. 10.1093/nar/gkt458

31. Heidarinia H, Tajbakhsh E, Rostamian M, Momtaz H (2023) Design and validation of a multi-epitope vaccine candidate against Acinetobacter baumannii using advanced computational methods. In Review

32. Sobolev OV, Afonine PV, Moriarty NW, et al (2020) A Global Ramachandran Score Identifies Protein Structures with Unlikely Stereochemistry. Structure 28:1249–1258.e2. 10.1016/j.str.2020.08.005

33. Desta IT, Porter KA, Xia B, et al (2020) Performance and Its Limits in Rigid Body Protein-Protein Docking. Structure 28:1071–1081.e3. 10.1016/j.str.2020.06.006

34. Kozakov D, Hall DR, Xia B, et al (2017) The ClusPro web server for protein–protein docking. Nat Protoc 12:255–278. 10.1038/nprot.2016.169

35. Guex N, Peitsch MC (1997) SWISS MODEL and the Swiss Pdb Viewer: An environment for comparative protein modeling. ELECTROPHORESIS 18:2714–2723. 10.1002/elps.1150181505

36. López-Blanco JR, Aliaga JI, Quintana-Ortí ES, Chacón P (2014) iMODS: internal coordinates normal mode analysis server. Nucleic Acids Res 42:W271–W276. 10.1093/nar/gku339

37. Craig DB, Dombkowski AA (2013) Disulfide by Design 2.0: a web-based tool for disulfide engineering in proteins. BMC Bioinformatics 14:346. 10.1186/1471-2105-14-346

38. Rice P, Longden I, Bleasby A (2000) EMBOSS: The European Molecular Biology Open Software Suite. Trends Genet 16:276–277. 10.1016/S0168-9525(00)02024-2

39. Grote A, Hiller K, Scheer M, et al (2005) JCat: a novel tool to adapt codon usage of a target gene to its potential expression host. Nucleic Acids Res 33:W526–W531. 10.1093/nar/gki376

40. Bibi S, Ullah I, Zhu B, et al (2021) In silico analysis of epitope-based vaccine candidate against tuberculosis using reverse vaccinology. Sci Rep 11:1249. 10.1038/s41598-020-80899-6

41. Walker, Morgan J, Nivens DK, Walker GM (2023) Characterization and Analysis of Mycobacteriophage Cain’s Genes

42. Laskowski RA (2001) PDBsum: summaries and analyses of PDB structures. Nucleic Acids Res 29:221–222. 10.1093/nar/29.1.221

43. Alhamadsheh MM, Musayev F, Komissarov AA, et al (2007) Alkyl-CoA Disulfides as Inhibitors and Mechanistic Probes for FabH Enzymes. Chem Biol 14:513–524. 10.1016/j.chembiol.2007.03.013

44. Dombkowski AA, Sultana KZ, Craig DB (2014) Protein disulfide engineering. FEBS Lett 588:206–212. 10.1016/j.febslet.2013.11.024

45. Yousaf M, Ullah A, Sarosh N, et al (2022) Design of Multi-Epitope Vaccine for Staphylococcus saprophyticus: Pan-Genome and Reverse Vaccinology Approach. Vaccines 10:1192. 10.3390/vaccines10081192

46. Weng, Chen (2011) Effect of Two-Step Fermentation by Rhizopus oligosporus and Bacillus subtilis on Protein of Fermented Soybean. Food Sci Technol Res 17:393–400. 10.3136/fstr.17.393

47. Tamang JP (2024) Unveiling kinema: blending tradition and science in the Himalayan fermented soya delicacy. J Ethn Foods 11:29

48. Sanjukta S, Sahoo D, Rai AK (2022) Fermentation of black soybean with Bacillus spp. for the production of kinema: changes in antioxidant potential on fermentation and gastrointestinal digestion. J Food Sci Technol 59:1353–1361. 10.1007/s13197-021-05144-y

49. Sharma, S, Padhi S, Kumari S, et al (2021) Bioactive compounds in fermented foods. In Bioactive compounds in fermented foods (pp. 48-69). CRC Press.

50. Choi YH, Cho SS, Simkhada JR, et al (2017) A novel multifunctional peptide oligomer of bacitracin with possible bioindustrial and therapeutic applications from a Korean food-source Bacillus strain. PLOS ONE 12:e0176971. 10.1371/journal.pone.0176971

51. Nguyen T, Nguyen CH (2020) Determination of factors affecting the protease content generated in fermented soybean by Bacillus subtilis 1423. Energy Rep 6:831–836. 10.1016/j.egyr.2019.11.011

52. Pahadi T (2022) EXTRACTION, PARTIAL PURIFICATION AND CHARACTERIZATION OF KINEMA PROTEASE PRODUCED FROM WHITE SOYABEAN

53. Liu Z, Pan J (2017) A practical method for extending the biuret assay to protein determination of corn-based products. Food Chem 224:289–293. 10.1016/j.foodchem.2016.12.084

54. Enechi O, Emilia CN (2013) A new colorimetric method for the determination of proteins. Adv Biol Res 7:159–162

55. Prasad S, Kashyap RS, Deopujari JY, et al (2006) Development of an in vitro model to study clot lysis activity of thrombolytic drugs. Thromb J 4:14. 10.1186/1477-9560-4-14

56. Weng Y, Yao J, Sparks S, Wang KY (2017) Nattokinase: an oral antithrombotic agent for the prevention of cardiovascular disease. Int J Mol Sci 18:523

57. Dabbagh F, Negahdaripour M, Berenjian A, et al (2014) Nattokinase: production and application. Appl Microbiol Biotechnol 98:9199–9206. 10.1007/s00253-014-6135-3

58. Peng Y, Yang X, Zhang Y (2005) Microbial fibrinolytic enzymes: an overview of source, production, properties, and thrombolytic activity in vivo. Appl Microbiol Biotechnol 69:126–132. 10.1007/s00253-005-0159-7

59. Stewart J, Manmathan G, Wilkinson P (2017) Primary prevention of cardiovascular disease: A review of contemporary guidance and literature. JRSM Cardiovasc Dis 6:204800401668721. 10.1177/2048004016687211

60. Ten Cate H (2021) Thrombosis: Grand Challenges Ahead! Frontiers Media SA

61. Gouda AM, Soltan MA, Abd-Elghany K, et al (2023) Integration of immunoinformatics and cheminformatics to design and evaluate a multitope vaccine against Klebsiella pneumoniae and Pseudomonas aeruginosa coinfection. Front Mol Biosci 10:1123411. 10.3389/fmolb.2023.1123411

62. Dorosti H, Eslami M, Negahdaripour M, et al (2019) Vaccinomics approach for developing multi-epitope peptide pneumococcal vaccine. J Biomol Struct Dyn 37:3524–3535. 10.1080/07391102.2018.1519460

63. Ismail S, Shahid F, Khan A, et al (2021) Pan-vaccinomics approach towards a universal vaccine candidate against WHO priority pathogens to address growing global antibiotic resistance. Comput Biol Med 136:104705. 10.1016/j.compbiomed.2021.104705

64. Lopéz-Blanco JR, Garzón JI, Chacón P (2011) iMod: multipurpose normal mode analysis in internal coordinates. Bioinformatics 27:2843–2850. 10.1093/bioinformatics/btr497

65. Ghosh P, Bhakta S, Bhattacharya M, et al (2021) A Novel Multi-Epitopic Peptide Vaccine Candidate Against Helicobacter pylori: In-Silico Identification, Design, Cloning and Validation Through Molecular Dynamics. Int J Pept Res Ther 27:1149–1166. 10.1007/s10989-020-10157-w

66. Yao X-Q, Skjærven L, Grant BJ (2016) Rapid Characterization of Allosteric Networks with Ensemble Normal Mode Analysis. J Phys Chem B 120:8276–8288. 10.1021/acs.jpcb.6b01991

