## Supplementary material for "*In Vitro* and Computational Evaluation of Thrombolytic Activity of Kinemakinase of Kinema, an Indigenous Fermented Food of Eastern Nepal": suplement Table 1

ORCHID ID:<https://orcid.org/0000-0003-3300-8191>

**Supplementary A**

**Physicochemical properties of kinema kinase**

**Table S1: Physicochemical and Immunological properties of Kinema kinase constructs**

| **Properties** | **Kinema kinase** |
| --- | --- |
| No. of amino acid | 381 |
| Molecular weight | 39507.48 Da |
| Theoretical PI | 9.04 |
| Instability Index | 28.84 |
| Positively Charged Amino acids | 30 |
| Negatively Charged Amino acids | 25 |
| Aliphatic Index | 81.73  (Thermally Stable) |
| GRAVY | -0.055 |
| Half-life in mammals | 30 hrs |
| Antigenicity | 0.7566 |
| Allergenicity | Allergen |
| Toxigenicity | Non-toxin |
| Solubility | 0.731 |
| Virulency | Virulent |
| Similarity with human/pig/mouse proteome | No |

**Table S2: Galaxyrefine models**

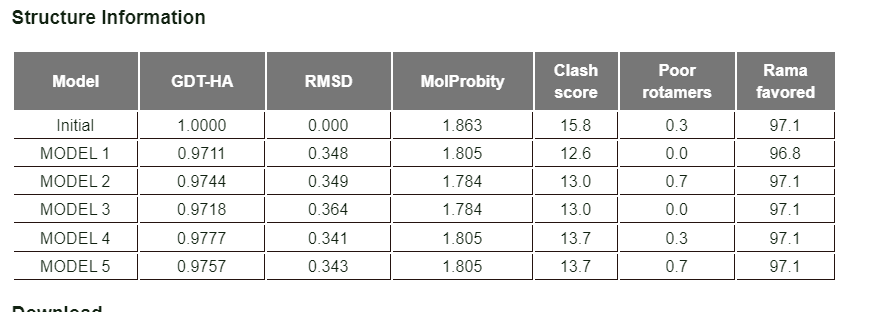

**Table S3: Top 5 Docked models**

| **Cluster** | **Members** | **Representative** | **Weighted Score** |
| --- | --- | --- | --- |
| **1** | 89 | Center | -985.3 |
|  |  | Lowest Energy | -1150.0 |
| **2** | 52 | Center | -934.1 |
|  |  | Lowest Energy | -1071.2 |
| **3** | 43 | Center | -1020.5 |
|  |  | Lowest Energy | -1125.8 |
| **4** | 38 | Center | -800.3 |
|  |  | Lowest Energy | -887.4 |
| **5** | 37 | Center | -830.0 |
|  |  | Lowest Energy | -951.7 |

Table S4: Selected pairs of amino acids for disulfide engineering

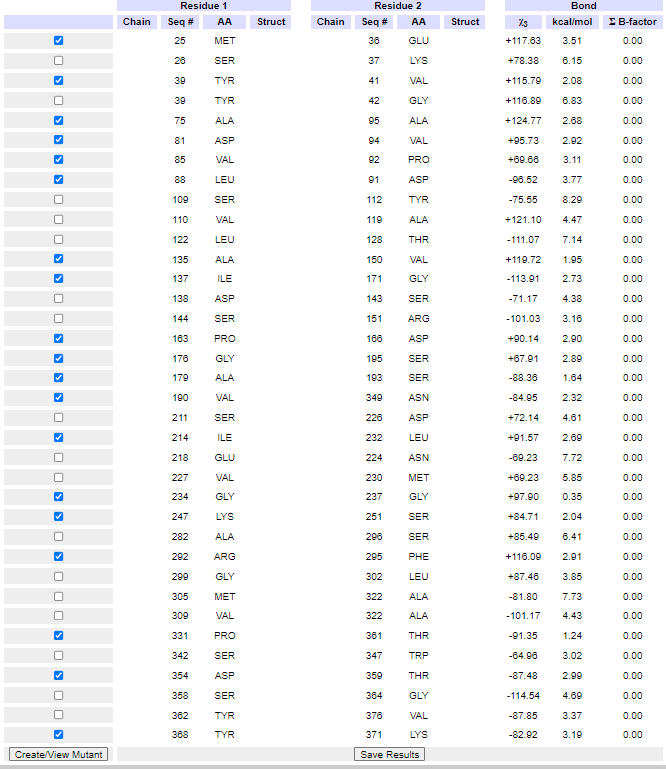
